# River biofilm bacteria as sentinels of national-scale freshwater ecosystems

**DOI:** 10.1101/2025.11.03.686311

**Authors:** Amy C. Thorpe, Susheel Bhanu Busi, Jonathan Warren, Lindsay K. Newbold, Joe D. Taylor, Kerry Walsh, Daniel S. Read

## Abstract

Freshwaters face increasing pressures from chemical, hydrological, and climatic changes, yet tools for assessing their condition remain limited. River biofilms, composed of diverse microbial communities, integrate environmental signals over space and time, making them sensitive indicators of river health. Using 16S rRNA gene sequencing of more than 1,600 biofilms collected across a national river network, we quantified bacterial diversity and community composition and applied network analysis to identify ecologically cohesive sub- communities with keystone taxa underpinning community stability. Alkalinity, dissolved oxygen, nitrate-nitrogen, and temperature were among the principal gradients shaping community composition. Threshold indicator analyses identified taxa with breakpoints along these gradients, revealing interpretable ecological thresholds. Our results demonstrate the potential for microbiome-based monitoring frameworks that complement existing biotic indices, enabling early detection of ecological changes and supporting the integration of genomic indicators into routine ecosystem assessment. This scalable approach offers a powerful strategy for managing freshwaters under accelerating anthropogenic pressures.

## Introduction

Freshwater underpins biodiversity, livelihoods, and climate resilience; however, freshwater ecosystems are among the most rapidly declining aspects of the biosphere (Dudgeon et al., 2007). A recent global assessment found that approximately one-quarter of assessed freshwater fauna are threatened with extinction, driven by pollution, water regulation and abstraction, invasive species, land use, and climate change (Sayer et al., 2025). These pressures interact nonlinearly across river networks and catchments, challenging conventional surveillance and policy frameworks. There is a growing consensus that next-generation biological evidence, particularly from genomic tools, should complement existing indicators to deliver earlier and more sensitive detection of ecosystem change and inform regulation (Andrei et al., 2025).

Microbial communities are central to this goal. They form the foundation of aquatic food webs (Clark et al., 2018), mediate core biogeochemical processes, including carbon and nutrient cycling (Falkowski et al., 2008), and respond rapidly to environmental variations at timescales relevant to management (Sagova-Mareckova et al., 2021). In rivers, benthic biofilms are complex assemblages of bacteria, archaea, fungi, and other microbial eukaryotes embedded within extracellular polymeric substances (EPS) that adheres to substrates such as stones (Battin et al., 2016; Besemer, 2015). In contrast to planktonic bacterial assemblages, which are often more transient and strongly influenced by water residence time (Read et al., 2015), biofilms form stable yet dynamically structured communities that reflect cumulative physiochemical conditions and watershed inputs over space and time (Brablcová et al., 2013). Recent studies have demonstrated that bacterial biofilm DNA can recover responses to land-use and pressure gradients, matching or even extending the diagnostic power of traditional biotic indices, highlighting the feasibility of incorporating bacterial communities into freshwater assessments (Hermans et al., 2024; Washington et al., 2013).

Despite these advances, the diversity and ecology of river biofilm bacterial communities, and the environmental drivers that structure them, are yet to be fully resolved at landscape and national scales (Veach et al., 2021). Studies across a range of scales have indicated strong filtering by water chemistry, temperature, and hydrology (Gautam et al., 2021, 2022; Lear et al., 2013). However, the relative importance of these gradients, their thresholds, and interactions remains poorly understood, particularly across large spatial scales. Addressing these gaps requires large, spatially balanced surveys coupled with robust multivariate and network approaches that can identify ecological breakpoints and candidate indicator taxa with clear mechanistic links to ecosystem change.

Here, we present the first national-scale characterisation of river biofilm bacterial communities in England, using 16S rRNA gene sequencing of 1,643 biofilm samples collected at 700 sites across the river network. We quantified bacterial diversity, community composition, and co-occurrence network structure, identified the principal physiochemical gradients associated with diversity, and used threshold indicator analyses to resolve taxa-specific breakpoints along these gradients. Together, these analyses provide a clear and interpretable microbial evidence base for next-generation freshwater biomonitoring and management.

## Results

### Bacterial community composition and diversity of river biofilms

Across the national-scale river network (Figure 1), bacterial biofilms displayed pronounced spatial and seasonal variability while maintaining a consistent taxonomic signature (Figure 2A). Pseudomonadota was the dominant phylum in 94.4% of samples, with a mean relative abundance of 0.50 (± 0.1 SD) across all samples and reaching a maximum relative abundance of 0.82. Other dominant members of the community included Bacteroidota (mean = 0.20 ± 0.1, max = 0.52) and Cyanobacteriota (mean = 0.12 ± 0.1, max = 0.87), confirming their importance in freshwater biofilms. Seasonal shifts were subtle, with Cyanobacteriota being slightly more abundant in spring, while other phyla showed little seasonal change.

**Figure 1.**
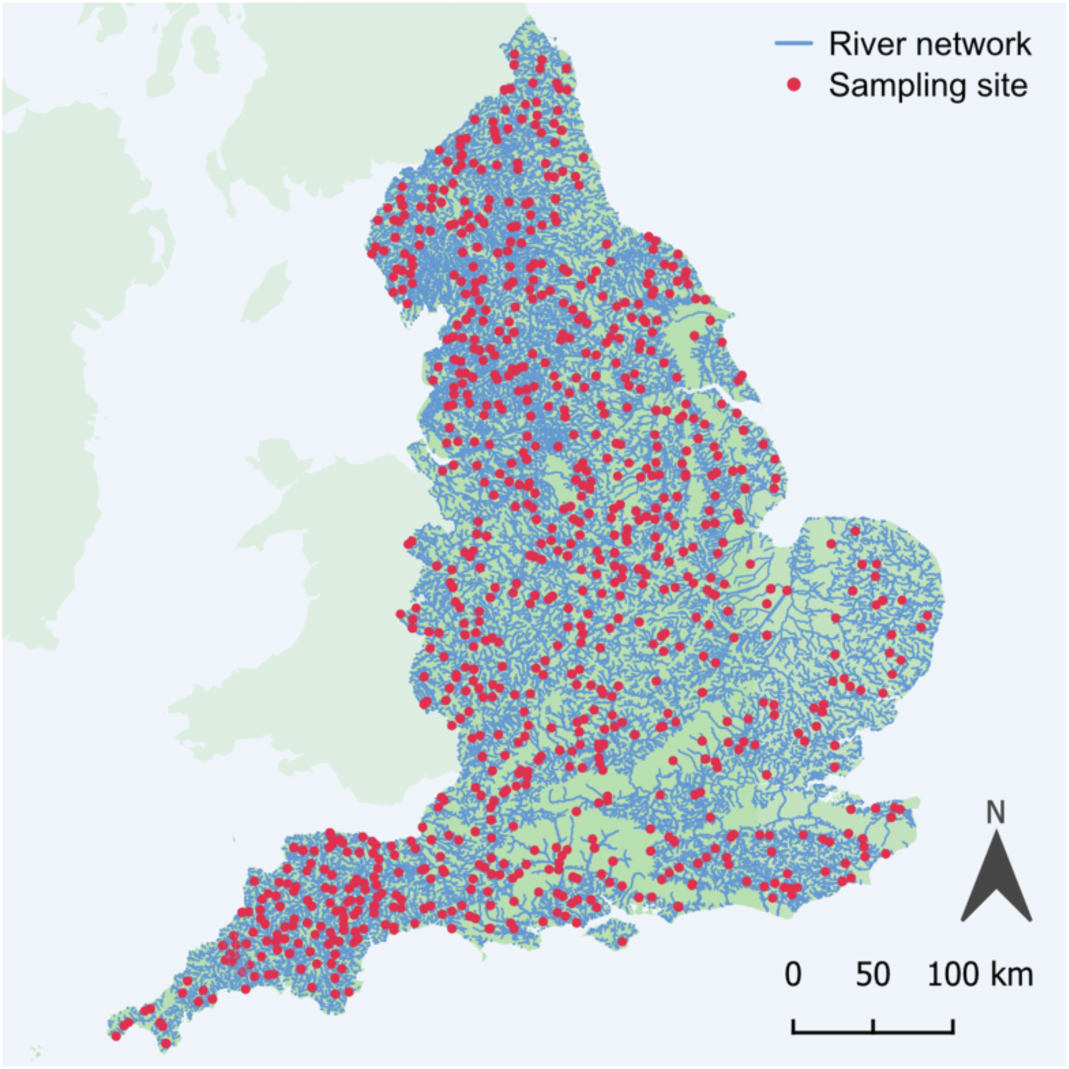
Biofilm sampling sites in rivers across England (contains OS data © Crown Copyright and database rights, 2025).

**Figure 2.**
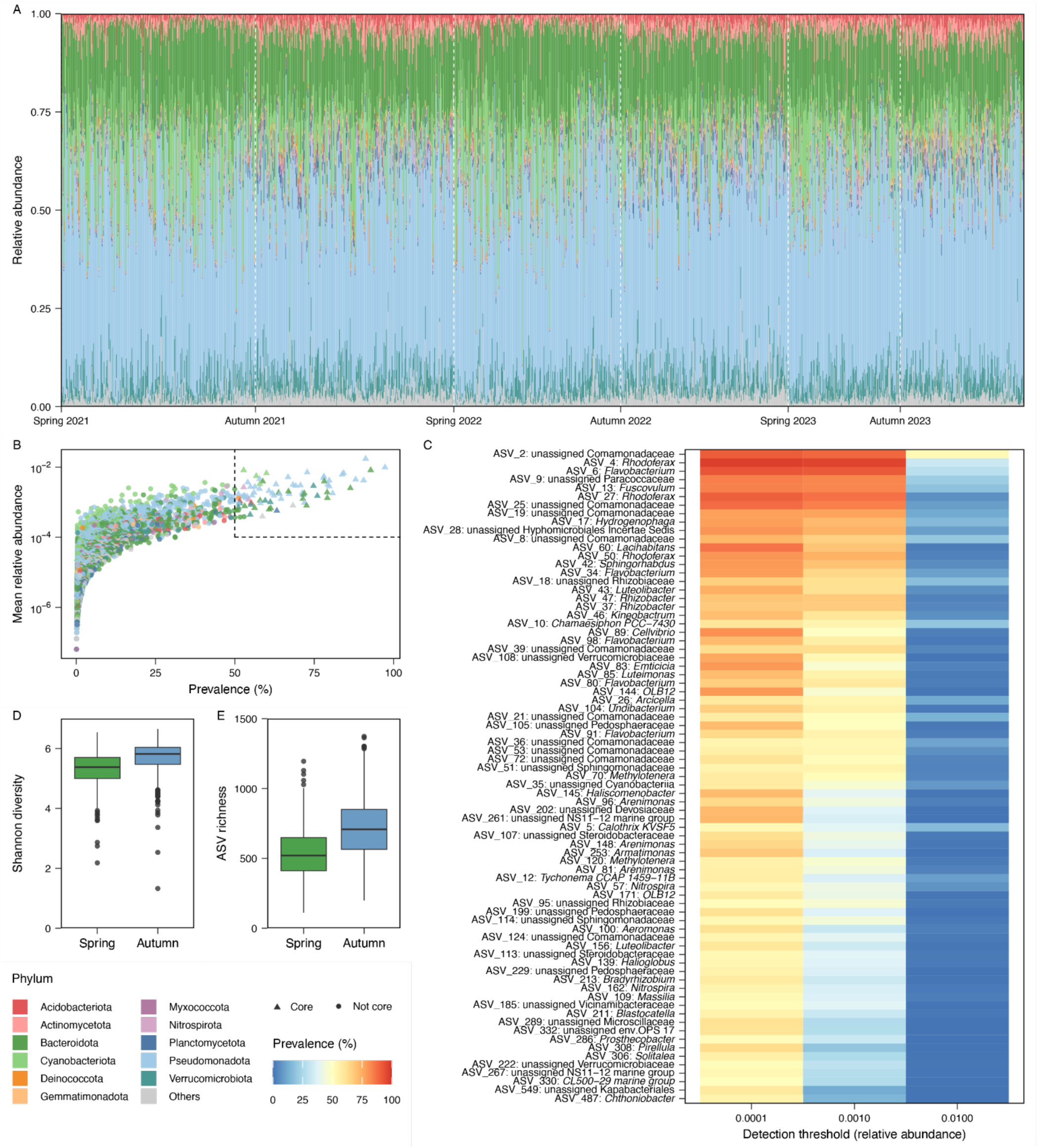
Bacterial community composition of river biofilms. (A) Relative abundance at the phylum level across seasons and years, where phyla with a mean relative abundance <0.005 are grouped as ‘Others’. (B) Mean relative abundance (log scale) and prevalence of ASVs, where triangles represent core ASVs. (C) Relative abundance and prevalence of the core community of ASVs, defined by a relative abundance >0.0001 in >50% of samples (approximated by the dashed box on panel B). (D) Shannon diversity and (E) ASV richness in spring and autumn.

The national-scale river biofilm community comprised many rare bacterial ASVs that exhibited relatively low prevalence and relative abundance (Figure 2B). Conversely, ASVs with a high prevalence tended to also show a higher relative abundance, and a relatively small core community of bacterial ASVs that were both abundant and prevalent across the river network (relative abundance >0.0001 in >50% of samples) was detected (Figures 2B and C). This included a total of 77 ASVs which encompassed a diversity of bacterial lineages. However, phylum-level patterns were evident, with Pseudomonadota and Bacteroidota containing numerous highly prevalent and abundant core ASVs (40 and 16 ASVs, respectively). Other members of the core community included Verrucomicrobiota (9 ASVs), Cyanobacteriota (4 ASVs), Acidobacteriota (2 ASVs), Nitrospirota (2 ASVs), Actinomycetota, Armatimonadota, Planctomycetota, and Candidatus Kapabacteria (each 1 ASV).

Alpha diversity and richness exhibited high levels of variation across the samples, with Shannon diversity ranging from 1.33 to 6.67, with a mean across all samples of 5.53 ± 0.6 (Figure 2D), and richness from 112 to 1,373, with a mean of 625.65 ± 210.21 (Figure 2E). Shannon diversity and richness exhibited some seasonal responses. Shannon diversity (spring mean = 5.32 ± 0.5 vs. autumn 5.72 ± 0.5; F_1,1552_ = 225.73, p <0.001; Figure 2B) and richness (spring mean = 536.71 ± 174.9 vs autumn 711.46 ± 205.8; F_1,1552_ = 327.85, p <0.001; Figure 2C) were both lower in spring than in autumn. Differences between sampling years were significant for Shannon diversity (F_2,1552_ = 12.99, p <0.001), but not for richness (F_2,1552_ = 2.10, p >0.05).

### Environmental drivers of river biofilm bacterial communities

NMDS based on beta diversity Bray-Curtis dissimilarities revealed a weak but consistent seasonal structure in community composition (R^2^ = 0.02, F = 32.06, p <0.001; Figure 3A), and only a marginal annual effect (R^2^ = 0.005, F = 3.99, p <0.001), suggesting that short-term temporal variability was limited compared to the environmental influences. Furthermore, the interaction between sampling year and season was not significant (R² = 0.002, F = 1.40, p >0.05), indicating that the seasonal signal in the community structure was consistent across years.

**Figure 3.**
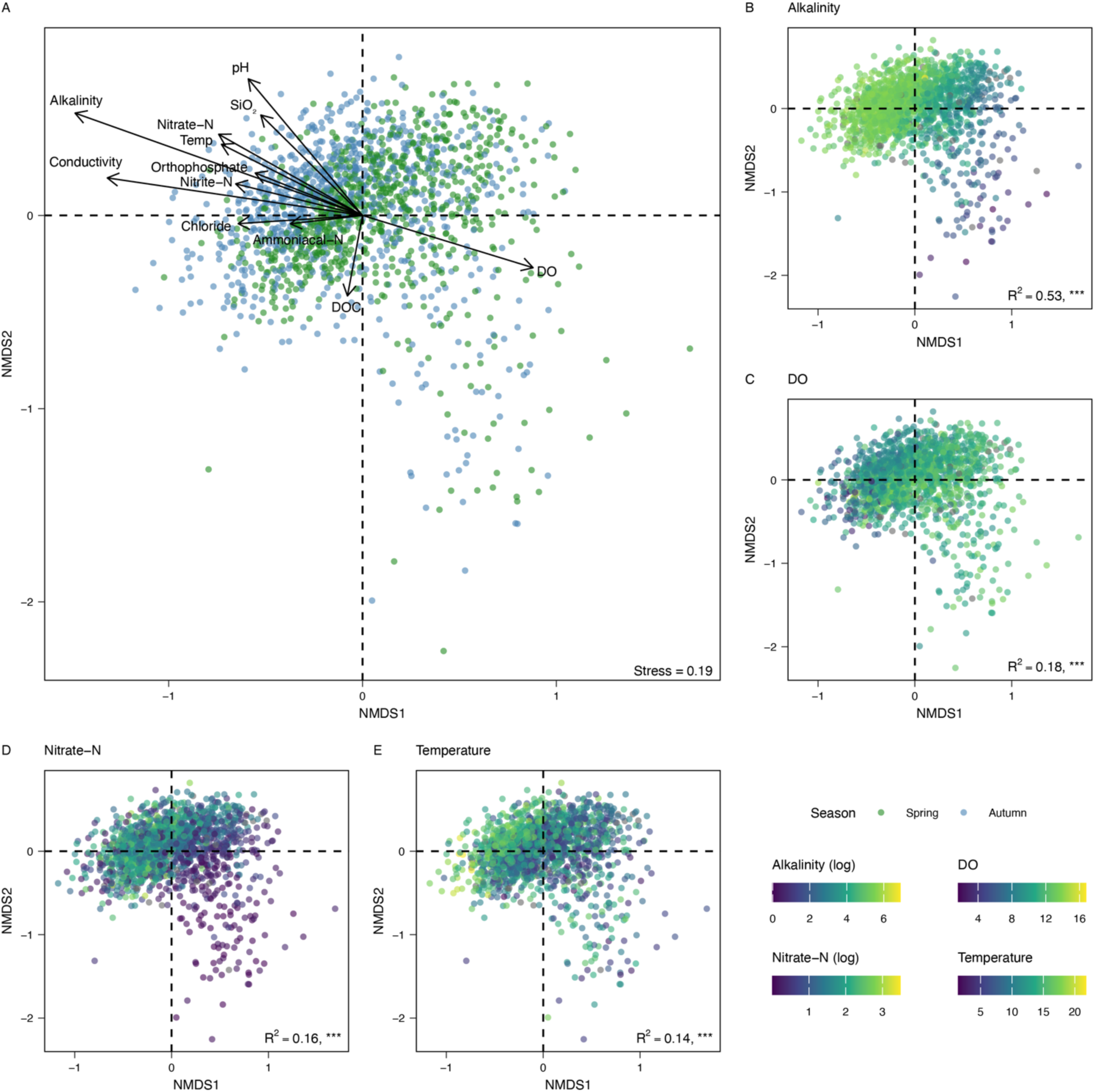
Non-metric multidimensional scaling (NMDS) of a beta diversity Bray-Curtis dissimilarity matrix of river biofilm bacterial communities. (A) NMDS, where samples are coloured by season. Vectors for water chemistry variables are fitted, and vector length is proportional to the strength of the correlation. NMDS where samples are coloured by gradients of selected water chemistry variables, (B) alkalinity, (C) DO, (D) nitrate-N, and (E) temperature. NA values are shown in grey. Alkalinity and nitrate-N were visualised on a logarithmic scale. Stress, R^2^, and significance levels are shown, p <0.001 = ***.

Among the measured variables, alkalinity exerted the strongest effect on biofilm community structure (R^2^ = 0.53, p <0.001; Figure 3B), followed by conductivity, pH, dissolved oxygen (DO), nitrate-nitrogen (nitrate-N), and temperature (R^2^ = 0.14-0.38, p <0.001; Figures 3B-E). Other measured factors contributed little to community structure (R^2^ <0.1; Supplementary Table 1).

### Bacterial indicators of freshwater ecosystem status

Co-occurrence network analysis revealed a highly modular community organisation (Figure 4A). The network, comprising 724 ASVs connected by 5,566 edges with an average degree of 15.38, had a modularity of 0.51 and a clustering coefficient of 0.46, reflecting a tendency of ASVs to form cohesive modules. These ASVs were organised into 20 ecological modules, with the three largest (M1, M2, and M3) dominated by Pseudomonadota, Bacteroidota, and Cyanobacteriota (Figure 4B), and containing several keystone taxa, defined as ASVs with high connectivity. Network heterogeneity (1.36) and centralisation (0.16) suggested moderate variation in connectivity and the presence of several highly connected taxa. Four ASVs were identified as connector nodes linking multiple modules (high among-module connectivity), all of which were in M1 or M3 and classified within the Pseudomonadota orders Rhodobacterales, Burkholderiales, and Sphingomonadales. Additionally, 21 ASVs were identified as module hubs, highly connected within their respective modules (high within-module connectivity), eight of which were also members of the core community. Module hubs were distributed across M1-M6 and included members of Pseudomonadota, Bacteroidota, Cyanobacteriota, Verrucomicrobiota, Nitrospirota, and Deinococcota (Figure 4C).

**Figure 4.**
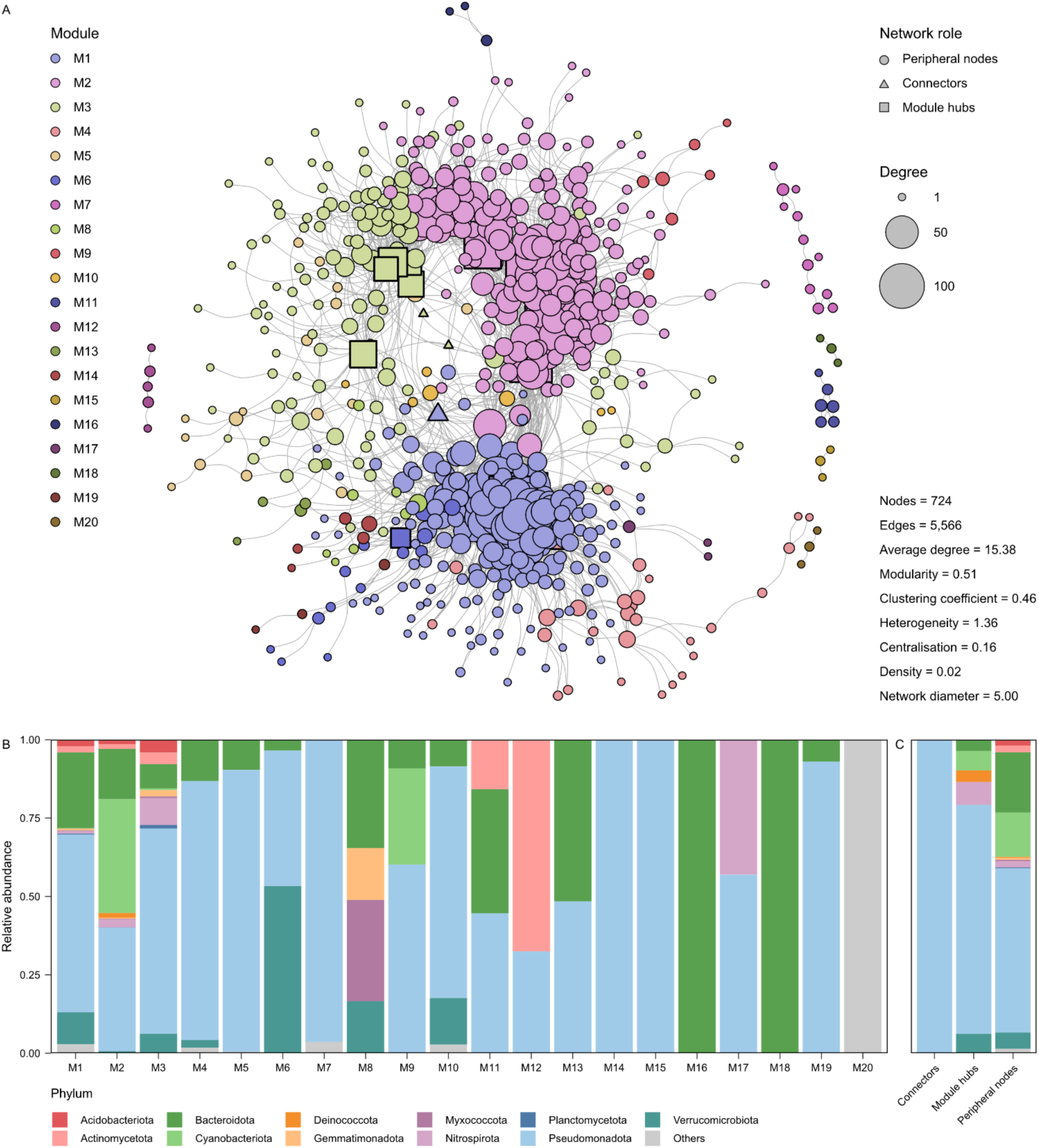
Co-occurrence network of river biofilm bacterial communities. (A) Co-occurrence network where nodes represent ASVs and are coloured by their module assignment. Node size is proportional to degree (number of connections) and node shape indicates network role where circles represent peripheral nodes, triangles represent connectors, and squares represent module hubs. Summary network metrics are shown. (B) Taxonomic composition of network modules, showing relative abundance of bacterial phyla within each module. (C) relative abundance of bacterial phyla by network role. Phyla with a mean relative abundance <0.005 are grouped as ‘Others’.

Several modules displayed strong correlations with key water chemistry variables, highlighting the environmental preferences among the modules. For instance, M1 and M3, which contained all the connector nodes and 12 module hubs, correlated most strongly with higher conductivity (M1 r = 0.45, p <0.001; M3 r = 0.54, p <0.001), alkalinity (M1 r = 0.45, p <0.001; M3 r = 0.53, p <0.001), ammonia-N (M1 r = 0.32, p <0.001; M3 r = 0.48, p <0001), nitrite-N (M1 r = 0.33, p <0.001; M3 r = 0.44, p <0.001), and orthophosphate (M1 r = 0.24, p <0.001; M3 r = 0.47, p <0.001). In contrast, M2, which contained seven module hubs, correlated with higher DO (r = 0.23, p <0.001) (Supplementary Figure 1).

Threshold indicator taxa analysis (TITAN) further identified community responses to key thresholds of environmental drivers: alkalinity, DO, temperature, and nitrate-N (Figures 5A- D). A total of 1,136 ASVs were identified as high-confidence and reliable indicators of alkalinity. Tolerant ASVs (those that respond positively to an increase in an environmental gradient) showed a change point at 122.5 mg L^-1^ of CaCO_3_ while sensitive ASVs (those that respond negatively to an increase in an environmental gradient) displayed a change point at 130.0 mg L^-1^ of CaCO_3_ (Figure 5A). A total of 911 ASVs were identified as indicators of DO, with tolerant ASVs showing a change point at 10.95 mg L^-1^ and sensitive ASVs at 9.73 mg L^-1^ (Figure 5B). Indicators of nitrate-N included 1,020 ASVs, with tolerant ASVs showing a change point at 4.07 mg L^-1^ and sensitive ASVs at 1.60 mg L^-1^ (Figure 5C). Finally, 911 indicator ASVs were identified for temperature, with tolerant ASVs showing a change point at 12.75 °C, and sensitive ASVs at 9.90 °C (Figure 5D).

**Figure 5.**
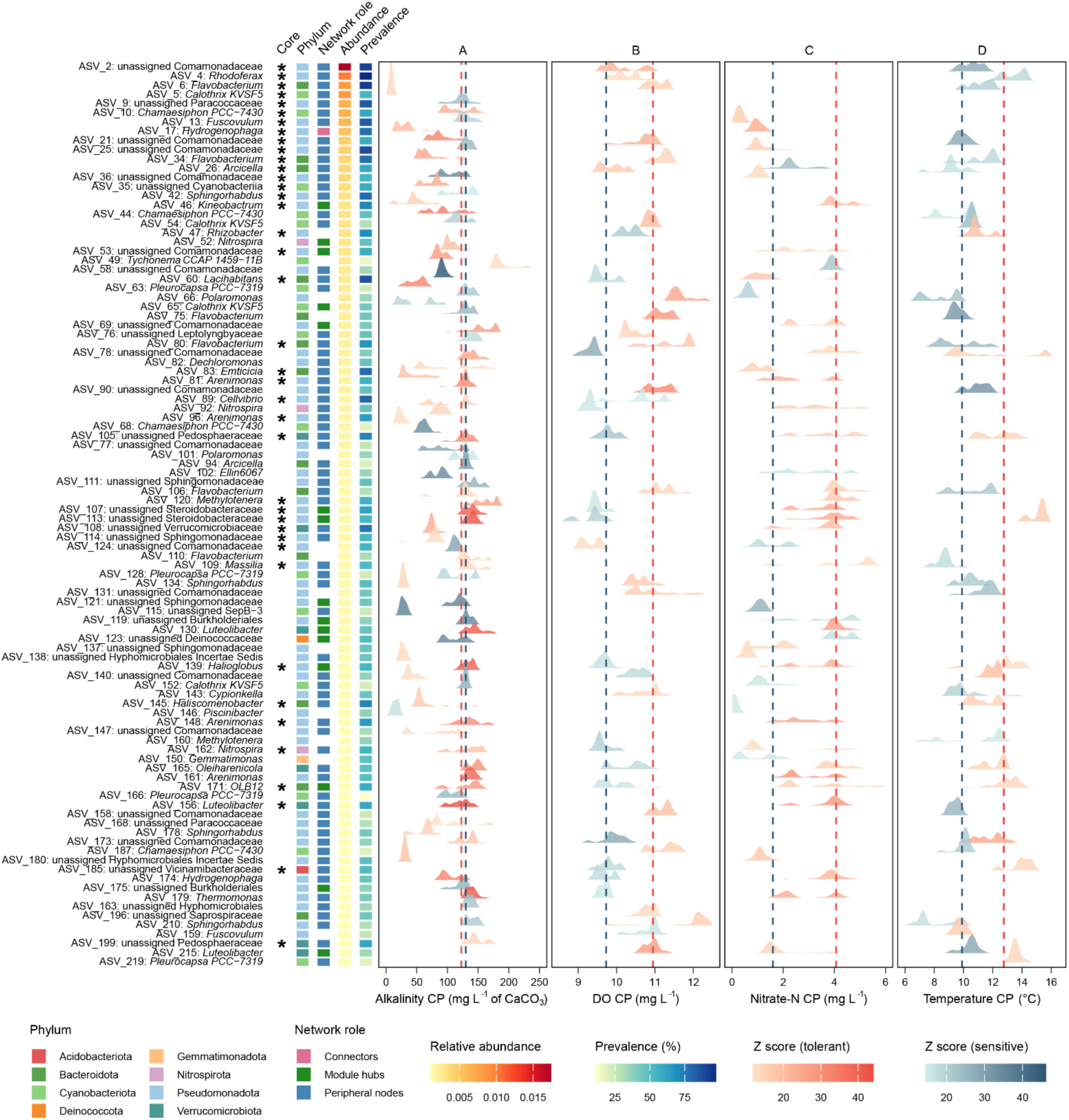
Threshold indicator taxa analysis (TITAN) of bacterial ASVs responding to (A) alkalinity, (B) DO, (C) nitrate-N, and (D) temperature. Ridges represent the distribution of bootstrap change points for each ASV, colour gradients correspond to the magnitude of the response at the median change point (Z score) for individual tolerant and sensitive ASVs, and dashed lines show the community-level change point (CP) for all tolerant (red) and sensitive (blue) ASVs. Core ASVs are marked with stars. The phylum, network role, mean relative abundance, and prevalence (%) are shown for each ASV. Each ASV is labelled by genus, and where the genus is unassigned, ASVs are labelled by the most specific assigned taxonomic level. Only ASVs with TITAN purity and reliability scores >0.95, Z scores >15, and mean relative abundance >0.001 are shown.

Many ASVs responded to multiple environmental gradients, with 578 (45.7%) serving as indicators of alkalinity, DO, nitrate-N, and temperature (Supplementary Figure 2). A smaller proportion of ASVs were uniquely identified as indicators of a single gradient. A total of 32 ASVs (2.5%) were exclusive to alkalinity, 10 ASVs (0.8%) to DO, 10 ASVs (0.8%) to nitrate- N, and 7 ASVs (0.6%) to temperature. Over half (55%) of the ASVs identified as indicator taxa by TITAN also belonged to the modules identified by co-occurrence network analysis, including all module hubs and connectors. Furthermore, 66 of the 77 core ASVs were module members and TITAN indicators.

## Discussion

Freshwater ecosystems are among the most biodiverse yet imperilled environments globally, with mounting evidence that chemical, hydrological, and climatic changes are driving widespread ecological degradation (Dudgeon et al., 2007; Reid et al., 2019). There is growing recognition that traditional biomonitoring approaches, while valuable, often fail to detect early or subtle changes in ecosystem conditions, particularly those induced by diffuse or cumulative pressures (Blackman et al., 2024). River microbial biofilms, which are complex, surface-attached multi-domain communities of microbes that integrate environmental signals over space and time, offer a promising but underutilised tool for freshwater ecosystem assessment (Battin et al., 2016). While benthic diatoms have been used as indicators of riverine ecological status (Taurozzi et al., 2024), bacterial communities remain largely unexplored in this context, despite being a dominant and functionally important component of biofilms with the potential to provide detailed, complementary insights into ecosystem condition.

This study represents the most comprehensive national-scale analysis of river biofilm bacterial communities to date, combining a spatially balanced sampling design with unified water chemistry and high-resolution 16S rRNA gene sequencing techniques. By analysing over 1,600 samples collected from 700 sites across England’s river network, we provided a detailed characterisation of benthic bacterial communities, identified the principal environmental gradients structuring their composition, and demonstrated the feasibility of using bacteria as threshold-based indicators of river conditions. Our findings provide a foundational evidence base for incorporating microbial metrics into freshwater biomonitoring frameworks, addressing longstanding calls to diversify the biological tools available for ecosystem surveillance (Andrei et al., 2025; Kuehne et al., 2023; Sagova-Mareckova et al., 2021).

Despite the pronounced spatial and environmental heterogeneity of river ecosystems across England, biofilm bacterial communities exhibited a consistent architecture dominated by Pseudomonadota, Bacteroidota, and Cyanobacteriota, alongside a diverse rare biosphere. This pattern mirrors metagenomic profiles from English rivers (Thorpe et al., 2025; Gweon et al., 2021) and global river systems (Lin et al., 2019; Busi et al., 2022). The persistence of a small core bacterial microbiome across England’s rivers highlights the dominance of generalist taxa adapted to prevailing physicochemical regimes, whereas rare and specialist lineages likely maintain functional redundancy (Reid et al., 2019; Shade et al., 2012). Seasonal patterns further reveal ecological succession, exemplified by lower diversity in spring, which may reflect a period of early biofilm development consistent with the microbial succession patterns reported in other temperate freshwater systems (Savio et al., 2015). This further suggests post-disturbance restructuring following high winter flows and light-driven shifts affecting phototrophic taxa, such as Cyanobacteriota (Paerl, 2017).

Multivariate analysis revealed that bacterial community composition was strongly associated with environmental gradients. Among all the variables tested, alkalinity explained the greatest proportion of beta diversity (R² = 0.53), followed by conductivity, pH, DO, nitrate-N, and temperature. These findings are consistent with previous continental and regional studies showing that carbonate buffering capacity and ionic strength are key determinants of bacterial community structure in lotic systems (Bier et al., 2023; Gweon et al., 2020; Niño-García et al., 2016; Ruiz-González et al., 2015). Importantly, these relationships were consistent across seasons and years, underscoring the robustness of these gradients as ecological filters at the national scale.

The pronounced influence of alkalinity likely reflects its association with pH and role in modulating the availability of dissolved inorganic carbon, both of which directly shape microbial physiology and metabolic potential (Newton et al., 2011; Pernthaler, 2017).

Similarly, nitrate-N and DO gradients likely reflect point and diffuse nutrient loading and organic pollution, as well as downstream oxidation processes. Temperature, although a weaker individual predictor, may interact with these variables to influence bacterial diversity and community composition, particularly among Cyanobacteriota, which showed seasonal enrichment in spring. However, it is important to recognise that water chemistry is also influenced by broader landscape-scale factors, such as underlying geology, hydrology, and land use, which reflect upstream catchment characteristics (Thorpe et al., 2025).

These observations align with emerging views that microbial assemblages respond to multivariate niche spaces rather than single stressors, necessitating analytical approaches capable of resolving complex nonlinear relationships (Widder et al., 2016). Our combination of ordination, network analysis, and threshold indicator modelling addresses this need by linking community structure to interpretable environmental breakpoints.

Network-based approaches have gained traction as tools for inferring microbial interactions and revealing community organisation beyond taxonomic profiles (Faust, 2021; Faust and Raes, 2012). Our analysis revealed that river biofilms are organised into connected, ecologically coherent modules, including sub-communities likely representative of shared environmental tolerances or functional guilds (Banerjee et al., 2018). The modularity and hub structure observed in our study are consistent with findings from soil and aquatic ecosystems, where tightly linked sub-communities often correspond to shared environmental tolerances or metabolic interdependencies (Banerjee et al., 2018; Layeghifard et al., 2017). Pseudomonadota dominated the module hubs and connectors, likely reflecting their metabolic versatility and ecological dominance in freshwater environments (Newton et al., 2011; Ruiz-González et al., 2015). Highly connected hub and connector taxa function as keystone species, maintaining within-module cohesion and cross-module interaction (Faust and Raes, 2012). In contrast, peripheral taxa with limited network links may act as specialists, confined to narrow environmental niches. Such modular partitioning balances community stability with flexibility, where hubs anchor resilience and specialists facilitate rapid adaptation to environmental change.

Threshold indicator analysis revealed distinct ecological breakpoints along gradients of alkalinity, nitrate-N, DO, and temperature, delineating transitions between tolerant and sensitive bacterial assemblages (Baker and King, 2010). These thresholds likely mark tipping points beyond which community composition, and potentially ecosystem function, becomes unstable. Notably, many core, hub, and connector ASVs were among the identified indicator taxa which exhibited strong responses to these gradients, thereby linking community stability and network topology to environmental sensitivity. This highlights their central role in shaping community dynamics and further supports the ecological coherence of these modules. Nearly half of the indicator taxa responded to multiple stressors, reflecting cross-sensitivity to the interacting drivers (Siddig et al., 2016). These taxa, which are central to network integrity and responsive to disturbance, represent promising indicators of ecosystem condition. Integrating network and threshold-based approaches strengthens the robustness and interoperability of microbial indicators, extending monitoring approaches already applied in soils and estuaries to riverine systems (Kelly et al., 2024; Nicolosi Gelis et al., 2024; Wang et al., 2013).

Collectively, these findings establish the potential for incorporating river biofilm bacterial communities into robust freshwater monitoring frameworks. By coupling a national-scale spatial survey design with DNA sequencing of river biofilms, we demonstrated that bacterial communities encode interpretable signals of freshwater environmental conditions, with links to water chemistry, modular organisation, and functional attributes. Importantly, our approach enables both community-level (i.e., community turnover and network structure) and taxon-level (i.e., indicator taxa and abundance thresholds) assessments of ecosystem change, providing multiple layers of diagnostic resolution. This flexibility is important for integration into regulatory frameworks, where different levels of detail may be required depending on the context, from rapid screening to detailed causal inference (Blackman et al., 2024; Sagova-Mareckova et al., 2021).

While further validation is required, particularly under different flow regimes, land uses, climatic contexts, and temporal scales, this study provides a blueprint for incorporating microbial metrics into catchment monitoring programmes. Genomic pipelines are scalable, reproducible, and amenable to automation, and align with the increasing digitisation of environmental surveillance (Kuehne et al., 2023). Moreover, the diagnostic power of the microbiome can be further enhanced by integrating functional analyses, trait-based modelling, and species sensitivity distributions coupled with predictive approaches to capture how microbial communities respond to environmental stressors. Future efforts should also aim to quantify the role of cross-domain and multi-trophic interactions, including between bacteria and phototrophic taxa, which may occur through resource exchange or EPS dynamics (Zancarini et al., 2017), as these relationships may strongly influence community stability and the sensitivity of microbial indicators. While network- and threshold-based approaches reveal associations within and between communities and the environment, they cannot fully resolve the mechanisms driving community assembly or the functional responses underpinning these patterns. Achieving a deeper mechanistic understanding will require integration with functional and experimental approaches, such as metagenomics and metatranscriptomics, alongside controlled manipulations, to test causal relationships between community structure, function, and ecosystem processes. These complementary approaches would extend insights from analyses of community assembly, helping to develop mechanistic, ecologically robust microbial indicators.

Our results reveal that biofilm bacterial communities are not only ecologically coherent but also diagnostically powerful. Freshwater biofilms are dominated by a core set of bacterial ASVs embedded within environmentally structured co-occurrence networks, and these ASVs are enriched with taxa that respond predictably to physicochemical drivers such as alkalinity, nitrate-N, DO, and temperature. We also identified hundreds of high-confidence indicator taxa, resolved community-level thresholds, and demonstrated temporal stability sufficient for routine monitoring. These attributes, combined with an analytically tractable approach, position biofilm bacteria as scalable sentinels of freshwater ecosystems.

By bridging community ecology, network science, and environmental genomics, this work advances the integration of microbial indicators into the management of freshwater ecosystems and offers a practical pathway to enhance the sensitivity, specificity, and explanatory power of river ecosystem assessments, supporting the transition to next-generation biomonitoring. As freshwater ecosystems face escalating pressures, tools that can detect changes earlier and explain them more clearly will be critical. The analysis of complex microbial communities contained within freshwater biofilms, as revealed in this study, is uniquely positioned to meet this challenge.

## Materials and Methods

### Sample collection

A total of 1,643 biofilm samples were collected from 700 sites across England (Figure 1). These sites form part of the Environment Agency’s River Surveillance Network that are routinely monitored to track changes in the health of England’s rivers. To ensure unbiased spatial coverage across the river network, sites were selected using a randomised and spatially balanced design (Brown et al., 2015). Sampling was conducted in spring (March to May) and autumn (September to November) over a three-year period from 2021 to 2023 to capture short-term temporal and seasonal variability. A total of 684 samples were collected in 2021 (339 in spring and 345 in autumn), 564 in 2022 (285 in spring and 279 in autumn), and 395 in 2023 (189 in spring and 206 in autumn). The majority of sites (594, 84.9%) were sampled twice over the three-year period (spring and autumn for one year), with a smaller proportion of sites (106, 15.1%) sampled once or three to seven times. Biofilm samples were collected from the river benthos at each site according to the standard protocol described in detail by Kelly et al. (2020). Briefly, benthic biofilms were collected by stone scraping and preserved in the field in 5 ml DNA preservation buffer (Warren et al., 2025). Samples were immediately transported to the Environment Agency (EA) National Laboratory at Starcross, Exeter, where they were concentrated by centrifugation and frozen before being transported on dry ice to the UK Centre for Ecology & Hydrology (UKCEH), Wallingford, where they were stored at -20 °C prior to DNA extraction.

### Environmental data collection

Surface water samples were collected from each sampling site to measure water chemistry variables, including temperature (°C), pH, alkalinity to pH 4.5 as CaCO_3_ (mg L^-1^), conductivity, and the concentration of dissolved oxygen (DO, mg L^-1^), dissolved organic carbon (DOC, mg L^-1^), orthophosphate (mg L^-1^), nitrate-nitrogen (nitrate-N, mg L^-1^), nitrite-nitrogen (nitrite-N, mg L^-1^), ammoniacal nitrogen (ammonia-N, mg L^-1^), and reactive silicon dioxide (SiO_2_, mg L^-1^). Water chemistry data can be accessed through the Water Quality Archive (Environment Agency, 2024). A mean was calculated for each variable using up to five independent measurements recorded over a 3-month period prior to and including the day of biofilm sampling.

### DNA extraction, PCR amplification, and sequencing

DNA was extracted from 100 µL of biofilm using the Quick-DNA Fecal/Soil Microbe Kit (Zymo Research, CA, U.S.) following a modified version of the manufacturer’s protocol to maximise DNA yield, as described by Newbold et al. (2025). A negative control with no sample added was included in every plate of 95 samples. The concentration and purity of the extracted DNA were checked using a NanoDrop 8000 spectrophotometer (Thermo Fisher Scientific, MA, U.S.), and the DNA concentration was further measured using the QuantiFluor ONE dsDNA kit (Promega, WI, U.S.). DNA was stored at 4 °C prior to PCR amplification.

A two-step PCR approach was used to first amplify the V4 region of the 16S rRNA gene with the forward, 515f-modified (5’-GTGYCAGCMGCCGCGGTAA-3’) and reverse, 806r- modified (5’-GGACTACNVGGGTWTCTAAT-3’) primers (Walters et al., 2016), and then uniquely barcode each sample in a second PCR using a series of forward and reverse index sequences to allow full demultiplexing of samples (Kozich et al., 2013). A detailed protocol, including reagents and thermocycling conditions for each PCR step, and purification, normalisation and pooling of PCR product, is described in Thorpe et al. (2024). Samples collected in 2021-22 and 2023 were sequenced across separate sequencing runs. Each 16S rRNA library was diluted to achieve a loading concentration of 1000 pM for paired-end sequencing on an Illumina NextSeq 2000 with a P1 flow cell and 30-40% PhiX control.

### Data processing

The 16S rRNA gene sequences were demultiplexed, and adapter sequences were trimmed using the Illumina FASTQ generation pipeline. Primer sequences were removed using Cutadapt v4.7 (Martin, 2011). The sequences were then processed using the DADA2 workflow (DADA2 R package v1.26.0) (Callahan et al., 2016). Quality distribution profiles were examined, and forward and reverse reads were truncated to 230 and 220 bp, respectively, to maintain a Q30 quality score across the reads. High-stringency filtering was performed with a maximum expected error of 2 and no ambiguous base pairs. Filtered reads were dereplicated into unique sequence variants and paired forward and reverse reads were aligned and merged with a minimum overlap of 12 bases. An amplicon sequence variant (ASV) abundance table was then constructed. The ASV tables generated from separate sequencing runs were merged, and chimeric sequences were removed. The naïve Bayesian classifier method (Wang et al., 2007) was implemented to assign taxonomy to each ASV using the SILVA v138.2 reference database (Quast et al., 2012) with a minimum bootstrap confidence of 80.

Non-bacterial ASVs (e.g., Eukaryota and Archaea, each accounting for 0.9% of ASVs, and chloroplast sequences, 2.1% of ASVs) and those unassigned at the phylum level (23.4% of ASVs) were removed. The data were further filtered to remove potential spurious ASVs with fewer than five reads and those present in fewer than five samples. Samples were then rarefied to a uniform sequencing depth of 10,000 reads, chosen based on the depth at which the richness plateaued for the majority of samples. Negative controls and 85 (5.2%) samples that did not meet this threshold were excluded from downstream analysis. A total of 24,067 bacterial ASVs from 1,558 samples were retained for the downstream analysis.

### Data analysis

Abundances were transformed into relative abundances and subsequently aggregated at the phylum level for visualisation using the phyloseq R package v1.51.0 (McMurdie and Holmes, 2013). The core community was identified using the microbiome R package v1.30.0 (Lahti and Shetty, 2017). An ASV was considered a member of the core community if it had a relative abundance >0.0001 in at least 50% of the samples (i.e., the prevalence threshold based on the proportion of samples in which the ASV was detected).

Alpha diversity measured as the Shannon index and richness at the ASV level were calculated using the vegan R package v2.6.10 (Oksanen et al., 2025). A two-way ANOVA was performed to determine whether there were significant differences in diversity and richness between seasons and years. Non-metric multidimensional scaling (NMDS) based on a Hellinger-transformed (Legendre and Gallagher, 2001) Bray-Curtis dissimilarity matrix of beta diversity was performed. Permutational multivariate analysis of variance (PERMANOVA) was conducted with 999 permutations using the adonis2() function to test for significant differences in beta diversity between the sampling seasons and years. To assess the relationship between environmental variables and beta diversity, water chemistry variables were fitted to the ordination space using the envfit() function with 999 permutations. Due to inter-correlations among related environmental parameters, such as orthophosphate and total phosphorus (TP), and nitrate-N and total oxidised nitrogen (TON), representative variables were used to reflect co-correlating gradients (Supplementary Figure 1).

ASVs were clustered into co-occurrence modules using the microeco (R package v1.14.0) network function. The network was computed with a relative abundance threshold of 0.0001, a prevalence threshold of 10% (1,249 ASVs), a Spearman correlation coefficient threshold of 0.40, and a p-value threshold of 0.01 (Liu et al., 2021). Modules were partitioned using the fast greedy modularity algorithm, and network roles were determined according to the within- and among-module connectivity scores. Connectors (nodes that are highly connected to other modules) were defined as ASVs with within-module connectivity ≤2.5 and among-module connectivity >0.62. Module hubs (nodes that are highly connected within their own module) were defined as ASVs with within-module connectivity >2.5 and among-module connectivity ≤0.62. Peripheral nodes (nodes with fewer connections, most of which are within their module) were defined as ASVs with within-module connectivity ≤2.5 and among-module connectivity <0.62 (Liu et al., 2021). Modules containing single pairs of ASVs were removed. The network was visualised using the igraph R package v2.1.4 (Csárdi and Nepusz, 2006). Spearman correlations were computed between module and phylum relative abundance, Shannon diversity, ASV richness and water chemistry variables.

Threshold indicator taxa analysis (TITAN) was performed using the TITAN2 R package v2.4.3 (Baker and King, 2010) to identify ASV and community-level thresholds along the water chemistry gradients. Using univariate regression tree partitioning and indicator analysis, TITAN separates the community into taxa that respond positively (tolerant) or negatively (sensitive) to environmental gradients. For each taxon, TITAN identifies the threshold (change point) at which the greatest change in relative abundance occurs, according to the magnitude of the response (Z score). Different taxa can have different change points, revealing complex and heterogeneous community dynamics. A community-level change point was also estimated, defined as the threshold at which the sum of individual Z scores was greatest, representing the point of greatest cumulative response of tolerant and sensitive taxa (Baker and King, 2010).

The environmental gradients tested were selected based on their strong association with beta diversity, as determined using envfit(). The selected gradients included alkalinity, DO, temperature, and nitrate-N. Although conductivity and pH are strong individual drivers, they were not investigated further because of their strong association with alkalinity (Supplementary Figure 1). These variables are closely related, where alkalinity reflects the buffering capacity of the water, which directly influences pH, and both are often linked with conductivity as a measure of ionic strength. Alkalinity was selected as a representative variable for this suite of interrelated gradients for downstream analysis to avoid redundancy and potential collinearity. The dataset was filtered to samples with complete observations (1,483 samples), and ASVs were filtered to include only those present in more than 10% of the samples (1,287 ASVs). TITAN was performed using 500 permutations for significance testing and 500 bootstrap replicates to estimate change point certainty. The results were filtered to include only ASV indicators identified as both pure (purity >0.95, i.e. consistent direction of response) and reliable (reliability >0.95, i.e. proportion of bootstrap permutations with a statistically significant response, p <0.05).

## Data availability

The R scripts to process and analyse the data are available at: https://www.github.com/amycthorpe/biofilm_16S_analysis. Raw sequence reads have been deposited in the European Nucleotide Archive (ENA) at EMBL-EBI under accession number PRJEB90117 available at: https://www.ebi.ac.uk/ena/browser/view/PRJEB90117. Sample accession codes and all the data presented are available on Zenodo at: https://doi.org/10.5281/zenodo.17515763.

## Author contributions

**ACT**: conceptualisation, methodology, data curation, investigation, formal analysis, visualisation, writing – original draft, writing – review & editing. **SBB**: conceptualisation, methodology, writing – original draft, writing – review & editing. **JW**: conceptualisation, funding acquisition, writing – review & editing. **LN**: conceptualisation, funding acquisition, methodology, writing – review & editing. **JT**: conceptualisation, funding acquisition, methodology, writing – review & editing. **KW**: conceptualisation, funding acquisition, writing – review & editing. **DSR**: conceptualisation, funding acquisition, methodology, writing – original draft, writing - review and editing.

## Acknowledgments

Thank you to Dr Jonathan Porter, Sean Butler, and Alan Wan from the Environment Agency National Laboratory Service for their support collating and shipping the biofilm samples for further processing. This work was funded by the Environment Agency under research project SC220034. ACT, SBB, and DSR were supported by Natural Environment Research Council (NERC) grant NE/X015947/1 and NE/Z000106/1. KW and JW were supported by NERC grant NE/X015777/1. The authors acknowledge the support of the Biotechnology and Biological Sciences Research Council (BBSRC), part of UK Research and Innovation; Earlham Institute Strategic Programme Grant Decoding Biodiversity BB/X011089/1 and its constituent work package - BBS/E/ER/230002C (Decode WP3 Linking Fine-Scale Microbial Diversity to Ecosystem Functions).

---

**Supplementary Figure 1.**
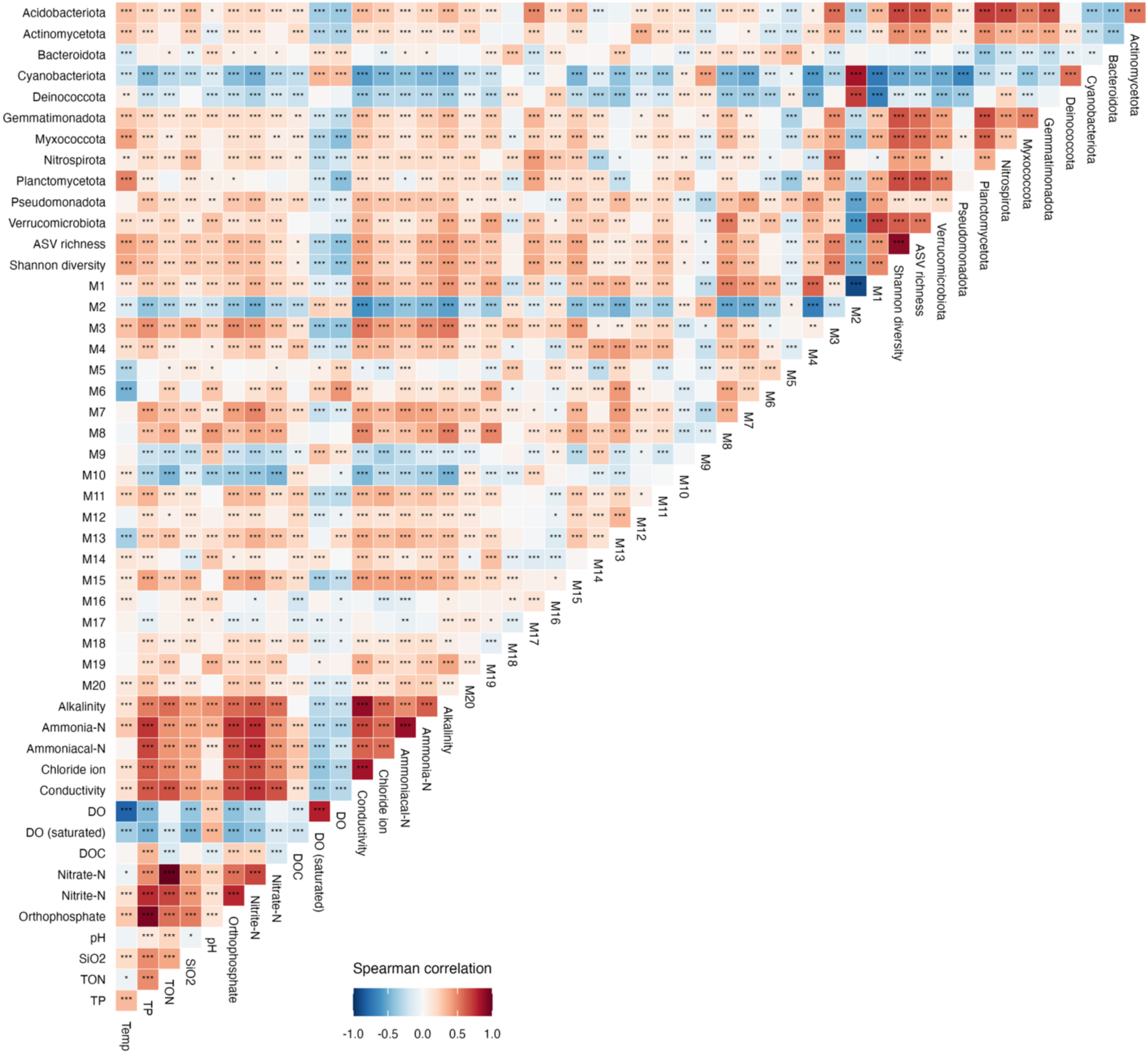
Spearman correlation matrix of the relative abundance of bacterial phyla, the relative abundance of co-occurrence modules, Shannon diversity and ASV richness, and water chemistry variables. The strength of the correlation is shown by the blue (more negative) to red (more positive) gradient. Significance levels are as follows: p <0.05 = *, p <0.01 =**, p <0.001 = ***.

**Supplementary Figure 2.**
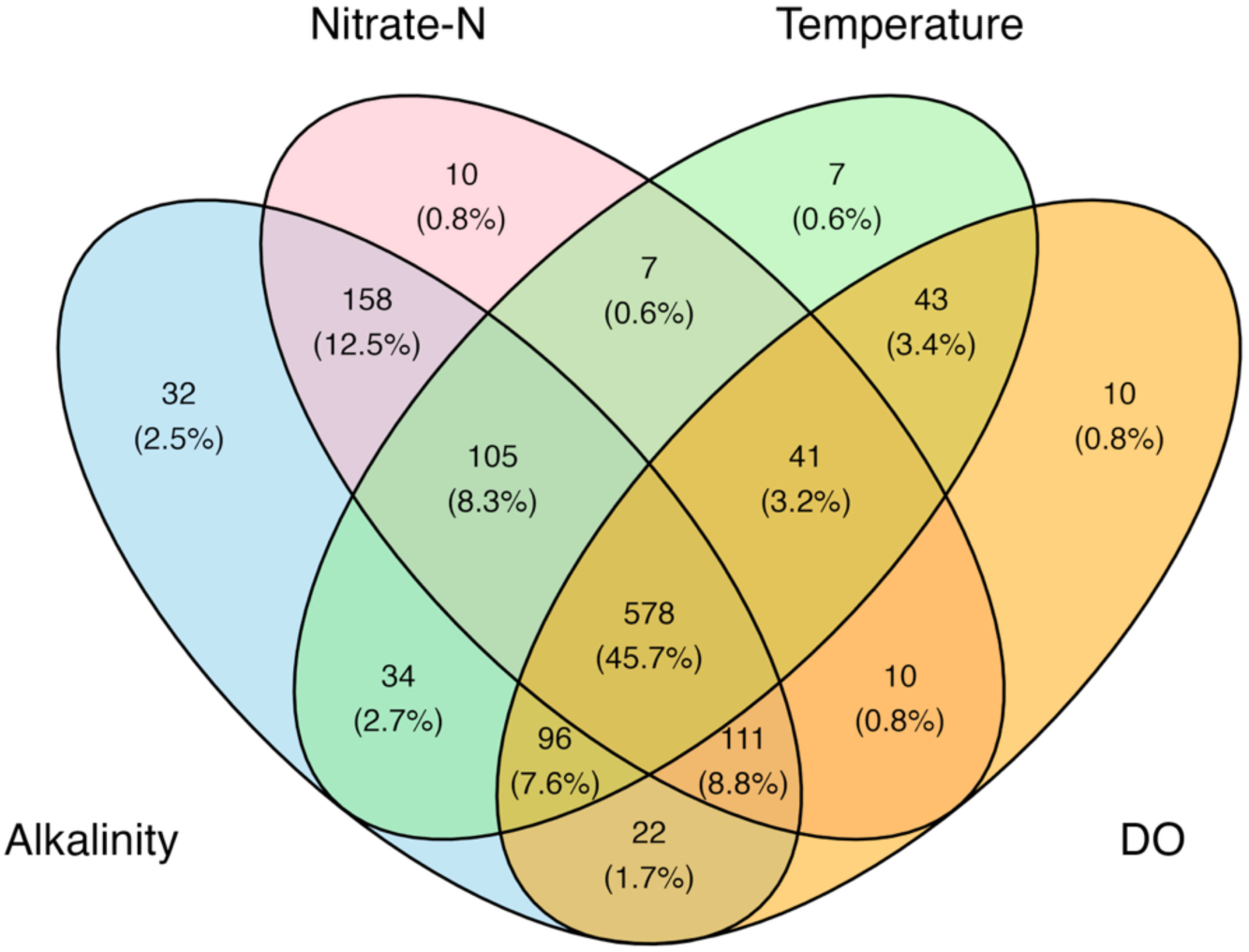
Number and proportion of shared and unique ASVs identified with TITAN as pure and reliable indicators of alkalinity, nitrate-N, temperature, and DO.

**Supplementary Table 1.** Results of the permutation test between the beta diversity Bray- Curtis dissimilarity matrix and water chemistry variables.

| Variable | R<sup>2</sup> | p |
| --- | --- | --- |
| Alkalinity | 0.53 | <0.001 |
| Conductivity | 0.38 | <0.001 |
| DO | 0.18 | <0.001 |
| pH | 0.18 | <0.001 |
| Nitrate-N | 0.16 | <0.001 |
| Temperature | 0.14 | <0.001 |
| SiO<sub>2</sub> | 0.12 | <0.001 |
| Nitrite-N | 0.10 | <0.001 |
| Chloride | 0.09 | <0.001 |
| Orthophosphate | 0.08 | <0.001 |
| DOC | 0.04 | <0.001 |
| Ammoniacal-N | 0.03 | <0.001 |

